# An oral inoculation infant rabbit model for *Shigella* infection

**DOI:** 10.1101/857086

**Authors:** Carole J. Kuehl, Jonathan D. D’Gama, Alyson R. Warr, Matthew K. Waldor

**Author notes:** equal contribution (Carole J. Kuehl, Jonathan D. D’Gama).

## Abstract

*Shigella* species cause diarrheal disease globally. Shigellosis is typically characterized by bloody stools and colitis with mucosal damage and is the leading bacterial cause of diarrheal death worldwide. Following oral ingestion, the pathogen invades and replicates within the colonic epithelium through mechanisms that rely on its type III secretion system (T3SS). Currently, oral infection-based small animal models to study the pathogenesis of shigellosis are lacking. Here, we found that oro-gastric inoculation of infant rabbits with *S. flexneri* resulted in diarrhea and colonic pathology resembling that found in human shigellosis. Fasting animals prior to *S. flexneri* inoculation increased the frequency of disease. The pathogen colonized the colon, where both luminal and intraepithelial foci were observed. The intraepithelial foci likely arise through *S. flexneri* spreading from cell-to-cell. Robust *S. flexneri* intestinal colonization, invasion of the colonic epithelium, and epithelial sloughing all required the T3SS as well as IcsA, a factor required for bacterial spreading and adhesion in vitro. Expression of the proinflammatory chemokine IL-8, detected with in situ mRNA labeling, was higher in animals infected with wild-type *S. flexneri* versus mutant strains deficient in *icsA* or T3SS, suggesting that epithelial invasion promotes expression of this chemokine. Collectively, our findings suggest that oral infection of infant rabbits offers a useful experimental model for studies of the pathogenesis of shigellosis and for testing of new therapeutics.

**Importance:** *Shigella* species are the leading bacterial cause of diarrheal death globally. The pathogen causes bacillary dysentery, a bloody diarrheal disease characterized by damage to the colonic mucosa and is usually spread through the fecal-oral route. Small animal models of shigellosis that rely on the oral route of infection are lacking. Here, we found that oro-gastric inoculation of infant rabbits with *S. flexneri* led to a diarrheal disease and colonic pathology reminiscent of human shigellosis. Diarrhea, intestinal colonization and pathology in this model were dependent on the *S. flexneri* type III secretion system and IcsA, canonical *Shigella* virulence factors. Thus, oral infection of infant rabbits offers a feasible model to study the pathogenesis of shigellosis and to develop and test new therapeutics.

## Introduction

*Shigella* species are Gram-negative, rod-shaped bacteria that cause bacillary dysentery, a severe and often bloody diarrheal disease characterized by inflammatory colitis that can be life-threatening (1). This enteric pathogen, which is spread by the fecal-oral route between humans, does not have an animal reservoir or vector (1). Annually, *Shigella* infections cause tens of millions of diarrhea cases and ∼200,000 deaths (2, 3). It is likely the leading cause of diarrheal mortality worldwide in individuals older than 5 years (2, 3). Most *Shigella* infections are attributable to *S. flexneri*, one of the four *Shigella* species, although in developed nations the prevalence of *S. sonnei* is higher (4–7).

The pathogen primarily causes colonic pathology that usually includes mucosal ulceration and erosion due to sloughing of epithelial cells, and is typically characterized by acute inflammation, with recruitment of neutrophils and plasma cells, congestion of blood vessels, distorted crypt architecture, and hemorrhage (8, 9). While inflammatory responses to *Shigella* invasion of colonic epithelial cells were thought to be the underlying cause of epithelial cell destruction and hemorrhage, recent evidence suggests that pathogen-mediated destruction of epithelial cells also plays a role in the development of pathology (10).

*Shigella* pathogenesis is attributable to a multifaceted set of virulence factors that enable the pathogen to invade and proliferate within the cytoplasm of colonic epithelial cells and evade host immune responses. The pathogen can also infect and rapidly kill macrophages (11). Most known virulence factors are encoded on a large (>200 kbp) virulence plasmid, which is required for *Shigella* pathogenicity (12–14). Key virulence determinants include a type III secretion system (T3SS) and its suite of protein effectors that are injected into host cells (15), and the cell surface protein IcsA, which directs polymerization of host actin and enables intracellular movement (16, 17). The force generated by intracellular actin-based motility allows the pathogen to form membrane protrusions into neighboring uninfected cells, which the pathogen subsequently enters. Cell-to-cell spread is thought to promote pathogen proliferation in the intestine and evasion of immune cells (11). The ∼30 T3SS effector proteins encoded by *Shigella* strains have varied functions, but primary roles include facilitating invasion of epithelial cells and suppression of host immune responses including cytokine production.

Among animals used to model infection, only non-human primates develop shigellosis from oral inoculation (18); however, the expense of this model limits its utility. Several small animal models of *Shigella* infection have been developed, yet none capture all the features of natural human infection. Historically, the Sereny test was used to identify *Shigella* virulence factors required for induction of an inflammatory response (19); however, this ocular model bears little resemblance to natural infection. The adult rabbit ligated ileal loop model has proven useful for the study of *Shigella* virulence factors (20). However, this model bypasses the normal route of infection and challenges the small intestine, which is not the primary site of pathology in human infections. Intra-rectal guinea pig infection induces colonic pathology and bloody diarrhea (21), and has been used to dissect the contribution of *Shigella* and host factors in several aspects of pathogenesis (22–24). Adult mice, the most genetically tractable mammalian model organism, are recalcitrant to developing disease when inoculated orally (25). As an alternative to oral inoculation, an adult mouse pulmonary model of *Shigella* infection involving intranasal inoculation of mice with *Shigella* has been developed (26); this model provides a platform to investigate host immune responses and vaccine candidates (27, 28), and has improved understanding of the innate immune response to *Shigella* infection (29). In contrast to adult mice, infant mice are susceptible to oral inoculation within a narrow window of time after birth, and inoculation with a high dose of the pathogen leads to mortality within a few hours; however, pathology is evident in the proximal small intestine rather than the distal small intestine or colon, and infected suckling mice do not develop diarrhea or intestinal fluid accumulation (30, 31). A zebrafish larvae model, in which the *Shigella* T3SS is required for pathogen virulence, has been useful for characterizing cell mediated innate immune responses to *Shigella* due to the ability to image infection in vivo (32, 33). Recently, an infant rabbit intra-rectal inoculation model was described in which animals develop disease and rectal pathology reminiscent of natural infections (10). The lack of a robust, oral inoculation-based, small animal model of shigellosis has limited understanding of the role of virulence factors in pathogenesis, particularly of the importance of such factors for enabling intestinal colonization and for generating pathology and clinical signs.

Here we found that oro-gastric inoculation of infant rabbits with *S. flexneri* results in severe disease resembling human shigellosis. Orally infected animals develop diarrhea and colonic pathology marked by damage to the epithelial cell layer and edema. Furthermore, the pathogen invaded and appeared to spread between colonic epithelial cells. We found that both the T3SS and IcsA were required for signs of disease, intestinal colonization and pathology. In addition, invasion of the pathogen into the epithelial cell layer was required for induction of host IL-8 expression. In situ mRNA labeling revealed that induction of IL-8 transcripts occurs primarily in cells adjacent to invaded epithelial cells, and not in the infected cells. Thus, our findings suggest that the oro-gastric infant rabbit model provides a powerful and accessible small animal model for further investigation of factors contributing to *Shigella* pathogenesis and for testing new therapeutics.

## Results

### Infant rabbits develop diarrhea following oro-gastric inoculation with *S. flexneri*

In previous work, we found that oro-gastric inoculation of infant rabbits with Enterohemorrhagic *Escherichia coli* (EHEC), *Vibrio cholerae*, and *V. parahaemolyticus* (34–36) leads to diarrheal diseases and pathologies that mimic their respective human counterparts. Here, we explored the suitability of oro-gastric inoculation of infant rabbits to model *Shigella* infection. *S. flexneri* 2a strain 2457T, a human isolate that is widely used in the research community as well as in challenge studies in humans (37), was used in this work. We utilized a streptomycin resistant derivative of this strain for infections to facilitate enumeration of pathogen colony forming units (cfu) in samples from the rabbit intestine. This strain, which contains a point mutation resulting in a K43R mutation in the small (30S) ribosomal subunit protein RpsL, retains the full virulence plasmid and grows as well as the parent strain.

In order to investigate infant rabbits as a potential *Shigella* host, we orally inoculated two to three-day-old rabbits that were co-housed with their dam and then monitored for signs of disease. There was considerable variability in the development of diarrhea and colonization in initial studies using suckling rabbits fed ad libitum. Previous work using four-week-old rabbits suggested that a milk component could protect animals from disease by degrading the *Shigella* T3SS components (38, 39); consequently, additional experiments were performed with infant rabbits separated from their lactating dam for 24 hours prior to inoculation. Using this protocol, we obtained more reliable clinical disease and robust intestinal colonization. By 36 hours post infection (hpi), the majority (59%) of animals developed diarrhea, which was grossly visible as liquid fecal material adhering to the fur of the hind region of the rabbits (Fig. 1A-C), and high levels of intestinal colonization (Fig. 2A); occasionally the diarrhea was frankly bloody. We chose the 36 hpi timepoint because from preliminary timecourse experiments we observed that all animals that were going to develop diarrhea developed disease by this timepoint and there was significant intestinal pathology at this time. Upon necropsy, the colon of infected animals was often bloody and contained liquid fecal material, in contrast to that of uninfected animals, which contained solid fecal pellets (Fig. 1D). Furthermore, some infected rabbits (27%) succumbed to infection rapidly and became moribund prior to 36 hpi, though not all of these animals developed diarrhea (Fig. 1A). Infected animals had highest bacterial burdens in the colon as well as the mid and distal small intestine (Fig 2A). The development of disease was associated with higher pathogen burdens in the colon (Fig. S1A). Separation of kits from the dam prior to inoculation led to a statistically significant elevation in intestinal colonization (Fig. S1B).

**Figure 1.**
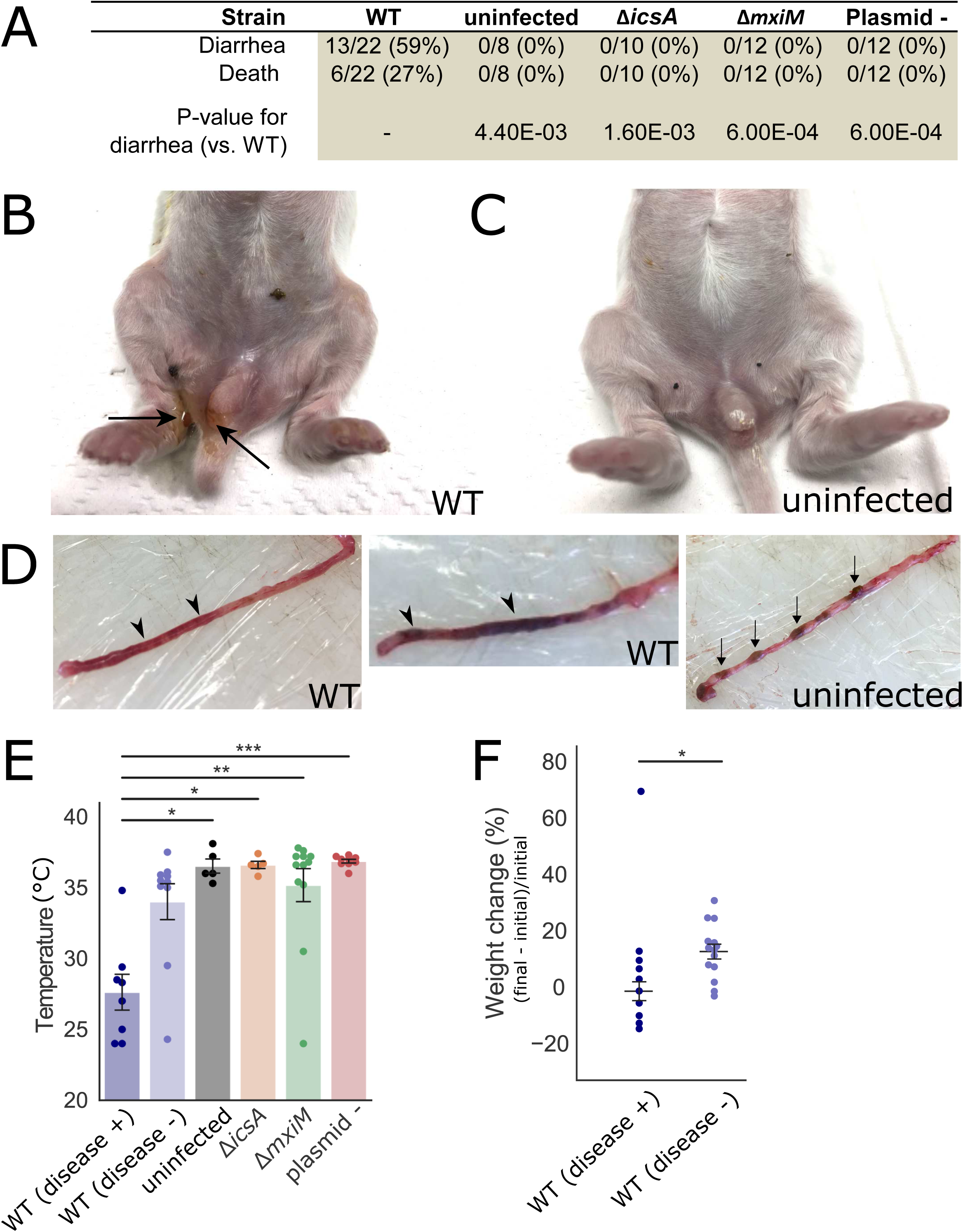
Clinical signs and gross pathology of infant rabbits following oro-gastric inoculation of *S. flexneri*. A. Clinical signs in infant rabbits infected with *S. flexneri* or isogenic mutant strains. Statistical significance for development of diarrhea between the animals in the WT group and in each of the other groups was determined using a Fisher’s exact test. B-C. Hind regions of animals inoculated with the WT strain (B) or an uninfected animal (C). Arrows indicate liquid feces stuck on anus and hind paws. D. Colons from animals inoculated with the WT strain (left) or of an uninfected animal (right). Arrowheads point to regions of liquid feces and arrows indicate solid fecal pellets. E. Body temperature of animals inoculated with the indicated strains 36 hpi or when they became moribund. Standard error of the mean values are superimposed. Disease +/− indicates whether or not animals developed diarrhea or became moribund early; all groups were compared to the WT (diarrhea +) group using a Kruskal-Wallis test with Dunn’s multiple-comparison. p-values: <0.05, *; <0.01, **; <0.001, ***. F. Percentage change in weight of infant rabbits infected with the WT strain, grouped by whether or not they developed disease (+/-). Percentage change in weight is calculated as difference between the final weight of the animal at 36 hpi or the last weight measurement taken when they became moribund (final) and the initial weight of the animal upon arrival in the animal facility (initial). Means and standard error of the mean values are superimposed. Groups were compared with a Mann-Whitney U test. p-values: <0.05, *.

**Figure 2.**
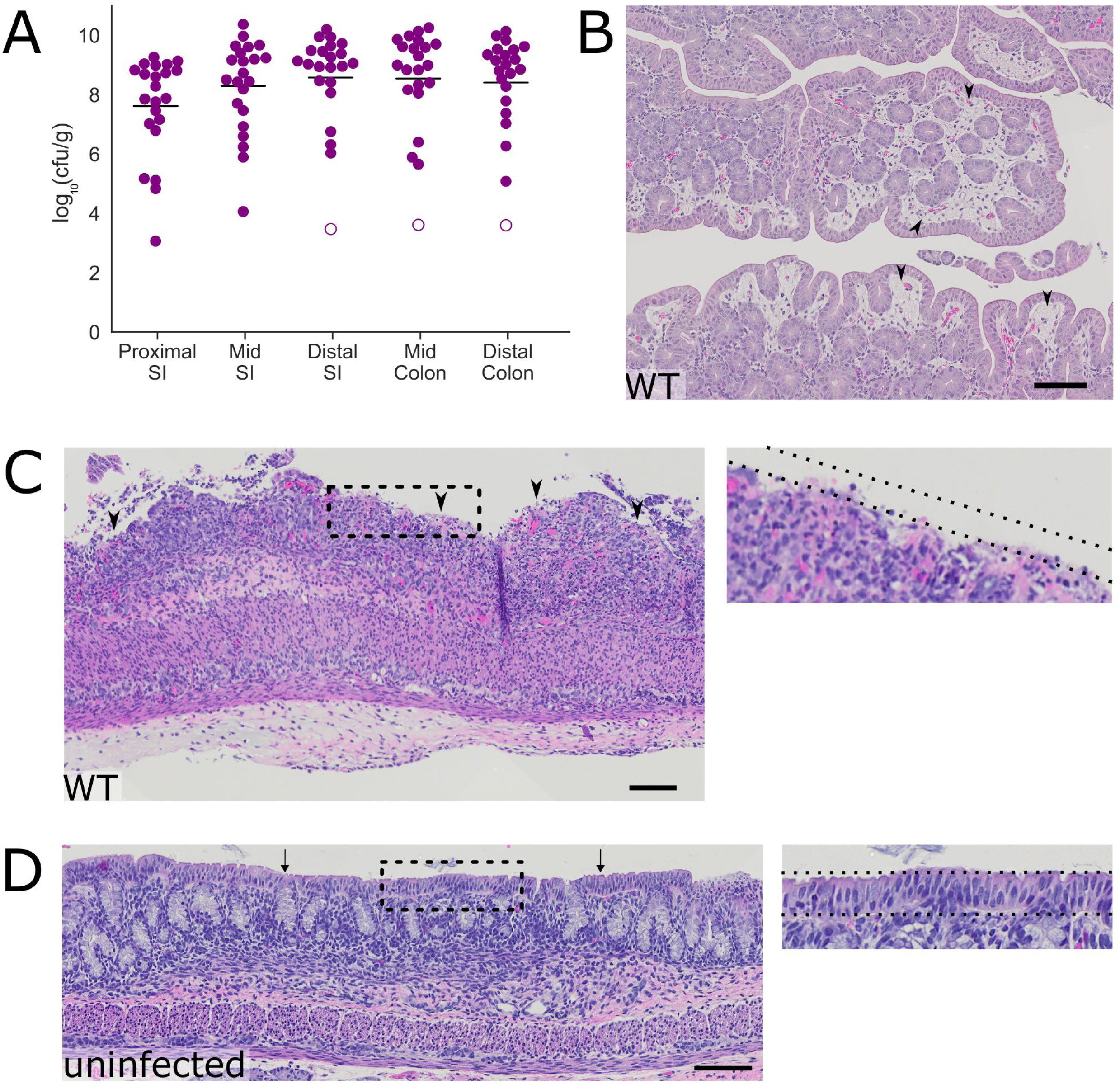
Intestinal colonization and colonic pathology in infant rabbits infected with *S. flexneri*. A. Bacterial burden of *S. flexneri* in the indicated intestinal sections 36 hpi; SI = small intestine. Each point represents measurement from one rabbit. Data plotted as log transformed colony forming units (cfu) per gram of tissue; mean values are indicated with bars. Open circles represent the limit of detection of the assay and are shown for animals where no cfu were recovered. B-D. Representative haematoxylin and eosin-stained colonic sections from infected animals (B,C) 36 hpi or uninfected animals (D). Arrowheads in (B) indicate areas of edema in the lamina propria. Arrowheads in (C) indicate areas where the epithelial cell layer is absent. Arrows in (D) point to the intact layer of epithelial cells seen in the colon. The dashed lines indicate the presence (inset, D) or absence (inset, C) of the epithelial cell layer. Scale bar is 100 μm.

Although not all *S. flexneri* inoculated animals developed signs of disease, infected rabbits that developed diarrhea or died early displayed additional disease signs. The animals that developed disease had significantly lower body temperatures than uninfected animals (8-9°C lower than uninfected, Fig. 1E), and they had significantly smaller gains in body weight than infected animals without disease (−2% vs 13%) (Fig. 1F) over the course of the experiment. Despite the relatively large intra- and inter-litter variation in body weight, with a constant pathogen dose per animal (1×10^9^ colony forming units, cfu), a lower initial body weight did not appear to be a risk factor for the development of disease (Fig. S1C).

Histopathologic examination of the intestines from infected rabbits revealed colonic pathology reminiscent of some of the features observed in infected human tissue, including substantial edema (Fig. 2B) as well as sloughing of colonic epithelial cells (Fig. 2C). In unusual cases, there was massive hemorrhage in the colonic tissue of infected rabbits (Fig. S2). Uninfected rabbits, which were similarly fasted, did not display colonic pathology and had no edema or disruption of the surface layer of epithelial cells (Fig. 2D). Notably, although the bacterial burden in the colon was similar to that of the distal small intestine (Fig. 2A), substantial pathology was not observed in the distal small intestine, suggesting organ-specific host factors influence the development of intestinal pathology.

### *S. flexneri* invades colonic epithelial cells after oro-gastric infection

Tissue sections from the colons of infected rabbits were examined with immunofluorescence microscopy to determine the spatial distribution of *S. flexneri* in this organ. The pathogen, which was labeled with an anti-*Shigella* antibody, was detected in the intestinal lumen and in many scattered foci within the epithelium (Fig. 3AB). At low magnification, the signal from the immunostained pathogen appeared to overlap with epithelial cells (Fig. 3A-C). At high magnification, immunostained *S. flexneri* was clearly evident within the boundaries of epithelial cells, which were visualized with phalloidin staining of actin and an antibody against E-cadherin (Fig. 3D). Several *S. flexneri* cells were frequently observed within an infected epithelial cell. In some infected epithelial cells, we observed *S. flexneri* cells associated with phalloidin stained actin tails (Fig. 3E), and in other foci, we observed *S. flexneri* in protrusions emanating from a primary infected cell with many cytosolic bacteria (Fig. 3F, asterisk), similar to structures seen in *Shigella* infections of tissue cultured cells (40, 41). The detection of actin tails and protrusions supports the hypothesis that the pathogen is actively spreading within the epithelial cell layer in the colon. *S. flexneri* cells were primarily localized to the epithelial cell layer and were infrequently observed in the lamina propria, the region of the intestinal wall directly below the epithelial cell layer. We did not find bacteria in the deeper layers of the intestine (Fig. 3A). Hence, following oro-gastric inoculation of infant rabbits with *S. flexneri*, the pathogen appears to proliferate both within the colonic lumen and in epithelial cells without penetration into deeper tissues.

**Figure 3.**
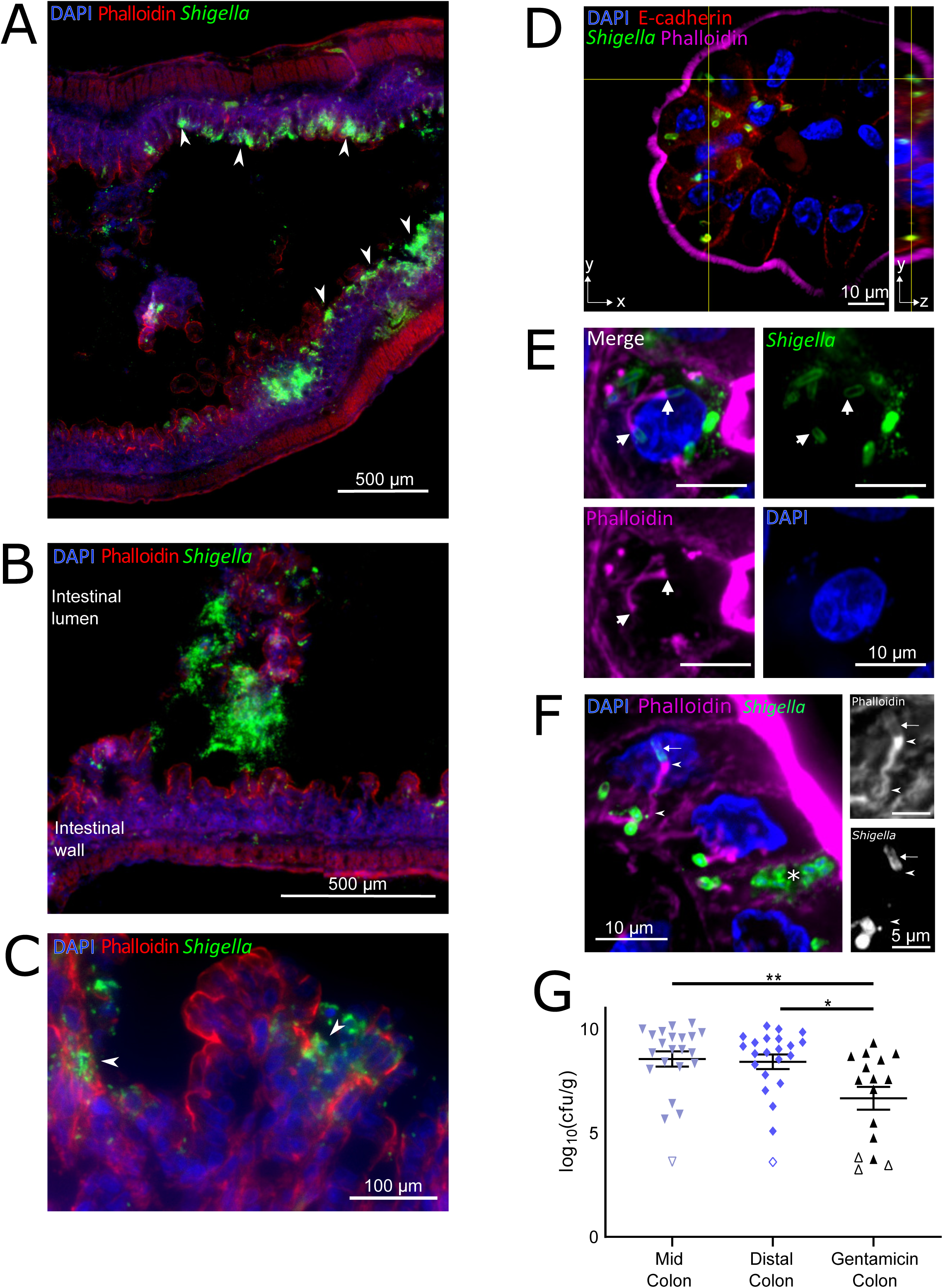
Localization of *S. flexneri* in the colons of infected infant rabbits. A-F. Immunofluorescence micrographs of *S. flexneri* in colonic tissue of infected rabbits 36 hpi. (A) *S. flexneri* bacteria were found in large numbers in epithelial foci (A, arrowheads point to selected foci). Scale bar is 500 μm. (B) *S. flexneri* bacteria in the lumen of the colon; the intestinal lumen and intestinal wall are labeled. Scale bar is 500 μm. (C) Arrowheads show infection foci where multiple neighboring cells contain intracellular *S. flexneri*. Scale bar is 100 μm. (D) Immunofluorescence z-stack micrograph of *S. flexneri* within colonic epithelial cells. Scale bar is 10 μm. Left (square) panel shows xy plane at a single z position, indicated by the horizontal axis of the cross-hairs in the yz projection. Right (rectangular) panel shows yz projection along the plane indicated by the vertical axis of the cross-hairs in the xy plane. Scale bar is 10 μm. (E) Immunofluorescence micrograph of *S. flexneri* associated with actin tails within colonic epithelial cells. Arrows point to poles of *S. flexneri* bacterial cells from which the actin tail is formed. Scale bar is 10 μm. (F) Immunofluorescence micrograph of *S. flexneri* forming protrusions during cell-to-cell spread between colonic epithelial cells. Asterisk marks a likely primary infected cell. Scale bar is 10 μm. Panels show zoomed region of phalloidin or anti-*Shigella* channels. Scale bar is 5 μm. Arrow points to actin surrounding the bacterial cell in a protrusion, arrowheads indicate the actin tail and actin cytoskeleton inside the protrusion at the pole of the bacterial cell and at the base of the protrusion. Blue, DAPI; green, FITC-conjugated anti-*Shigella* antibody; red (A-C or magenta in D-F), phalloidin-Alexa 568; and when present, red (D), anti-E-cadherin. G. Bacterial burden of *S. flexneri* WT strain in the indicated intestinal sections 36 hpi. Each point represents measurement from one rabbit. Data plotted as log transformed colony forming units (cfu) per gram of tissue; means and standard error of the mean values are superimposed. Open symbols represent the limit of detection of the assay and are shown for animals where no cfu were recovered. Statistical significance was determined with a Kruskal-Wallis test with Dunn’s multiple comparison. p-values: <0.05, *; <0.01, **.

We also measured the burden of intracellular *S. flexneri* in the colon using a modified gentamicin protection assay previously used to study the intracellular burden of *Listeria monocytogenes* and *Salmonella enterica* serovar Typhimurium in murine intestinal tissues (42–46). After dissecting intestines from infected infant rabbits, colonic tissue was incubated with gentamicin, an antibiotic that selectively kills extracellular (i.e. luminal) bacteria. We observed an ∼2 log decrease in bacterial burden after gentamicin treatment (Fig. 3G), suggesting that only a small portion of *S. flexneri* in the colon are intracellular.

### IL-8 transcripts are often observed in epithelial cells near infected cells

We next investigated aspects of the infant rabbit host innate immune response to *S. flexneri* infection. IL-8, a proinflammatory CXC family chemokine that recruits neutrophils (47), has been shown to be elevated during *Shigella* infection in animal models (10, 21, 48) and in humans (49, 50). However, in preliminary experiments it was difficult to detect significant elevations of IL-8 transcripts in bulk colonic tissue using a qPCR-based assay. Due to the patchiness of the infection foci observed through immunofluorescence imaging of colonic tissue, we wondered whether a localized response to infection might be masked when analyzing bulk intestinal tissue specimens. Local expression of IL-8 mRNA in *S. flexneri*-infected tissue was assessed using RNAscope technology, a sensitive, high-resolution in situ mRNA imaging platform that permits spatial analysis of mRNA expression. In the colon, we detected localized expression of IL-8 mRNA in colonic epithelial cells near infection foci (Fig. 4AB). In contrast, very few IL-8 transcripts were detected in the colons of uninfected kits (Fig. 4CD). Combined detection of IL-8 and *S. flexneri* demonstrated that IL-8 expressing cells were typically near cells containing *S. flexneri*, but not themselves infected with the pathogen (Fig. 4A, B & D, & S3). The majority (>90%) of infected epithelial cells did not express IL-8 mRNA, while >40% of these infected cells were adjacent to uninfected cells that did express IL-8 mRNA. Several T3SS effectors from *S. flexneri*, e.g. IpgD (51) and OspF (52), have been shown to reduce IL-8 expression in infected cells, which may explain the weak or absent IL-8 production in infected cells. There was a wide range in the prevalence of IL-8 producing cells in infected animals (Fig. 4D & S3). The variability of IL-8 expression after infection may reflect the patchiness of *S. flexneri* invasion along the colon (Fig. 3A). Together, these observations suggest that *S. flexneri* infection induces IL-8 mRNA expression (and perhaps additional cytokines as well) in infant rabbits.

**Figure 4.**
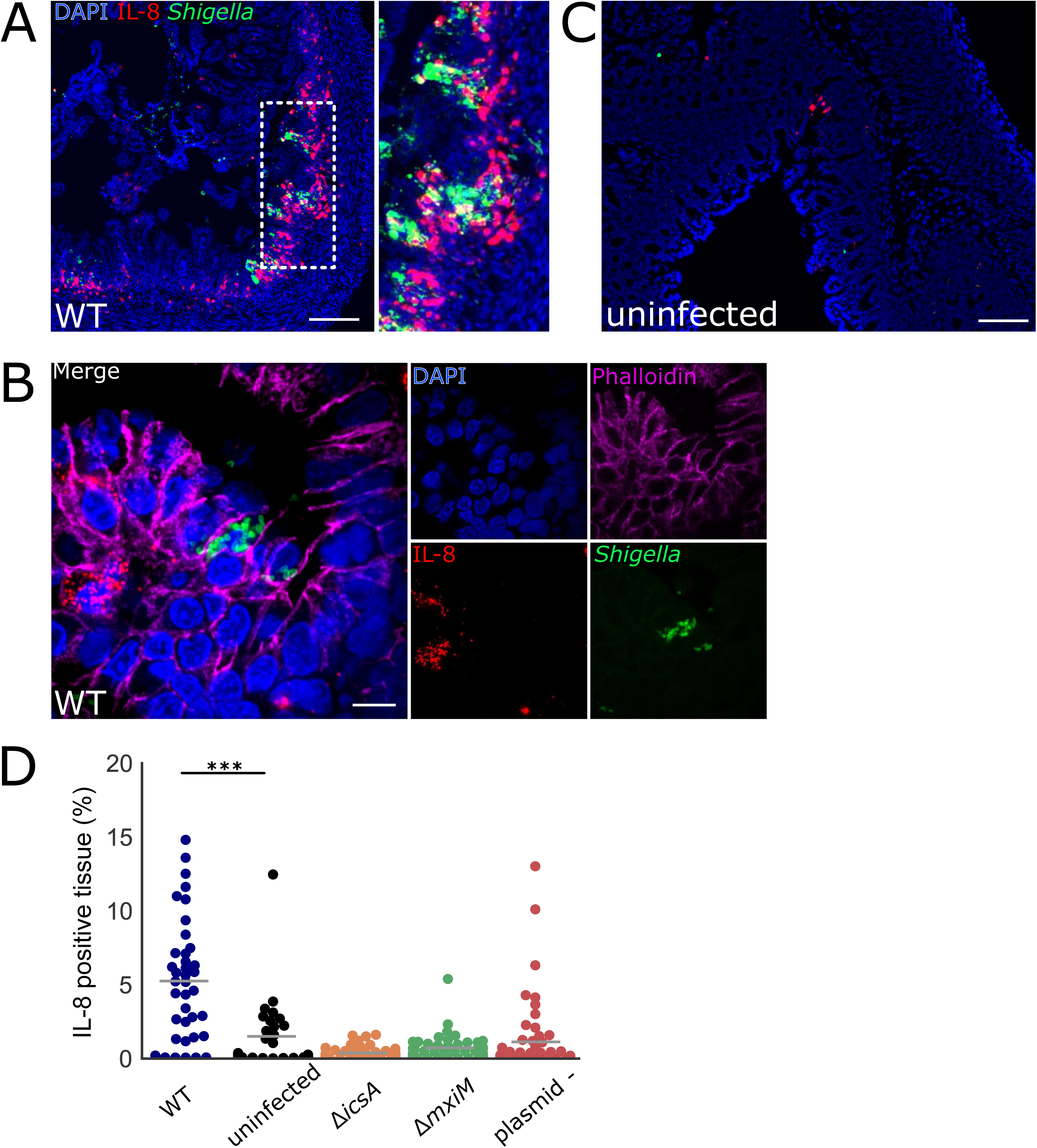
Colonic IL-8 mRNA in rabbits infected with *S. flexneri*. A-C. Immunofluorescence micrographs of colonic sections from infant rabbits infected with WT *S. flexneri* (A, B) or uninfected control (C). Sections were stained with an RNAscope probe to rabbit IL-8 (red), an antibody to *Shigella* (FITC-conjugated anti-*Shigella* green), and with DAPI (blue). (A) Colon section infected with WT *S. flexneri.* Inset on right of A depicts magnified view of boxed area on left image. Scale bar is 200 μm. (B) High magnification of colonic epithelium infected with WT *S. flexneri.* Sections were also stained with anti-E-cadherin antibody (magenta). Scale bar is 10 μm. (C) Uninfected colon section. Scale bar is 100 μm. D. Percentage of IL-8 expressing cells in each field of view from colonic tissue sections stained with probe to rabbit IL-8 from rabbits infected with the indicated strain. See methods for additional information regarding the determination of these measurements. Mean values are indicated with bars. All groups were compared to the sections from the uninfected animals. Statistical significance was determined using a Kruskall-Wallis test with Dunn’s multiple comparison. P-values: <0.001, ***.

### Narrow bottleneck to *Shigella* infection of the infant rabbit colon

We attempted to use transposon-insertion sequencing (TIS) to identify genetic loci contributing to *S. flexneri* colonization and pathogenesis, as we have done with *V. cholerae* (53, 54), *V. parahaemolyticus* (55), and EHEC (56). Initially, a high-density transposon mutant library in *S. flexneri* was created using a *mariner*-based transposon that inserts at TA dinucleotide sites in the genome. The library included insertions across the entirety of the genome, including the virulence plasmid (Table S1). Infant rabbits were inoculated with the transposon library and transposon mutants that persisted after 36 hpi were recovered from the colon. Comparison of the frequencies of insertions in the input and output libraries revealed that the output transposon libraries recovered from rabbit colons only contained ∼20% of the transposon mutants that were present in the input library. These observations suggest that there is a very narrow bottleneck for *S. flexneri* infection in rabbits, leading to large, random losses of diversity in the input library. These random losses of mutants confound interpretation of these experiments and precludes accurate identification of genes subject to in vivo selection. Modifications to the in vivo TIS protocol will be necessary to apply TIS to identify additional *S. flexneri* colonization factors.

### Canonical *S. flexneri* virulence factors are required for intestinal colonization and pathogenesis

Next, we investigated the requirement for canonical *Shigella* virulence factors in intestinal colonization and disease pathogenesis. First, we tested a strain that lacked the entire virulence plasmid, which contains most of the known virulence factors encoded in the *S. flexneri* genome, including the T3SS. As anticipated, this strain was avirulent; animals inoculated with the plasmid-less (plasmid -) *S. flexneri* strain did not die or develop diarrhea or reduced temperature (Fig. 1A & E). We also tested isogenic mutants that lack one of two key virulence factors: IcsA (Δ*icsA* strain), which is required for intracellular actin-based motility and cell-to-cell spreading, and MxiM (Δ*mxiM* strain), which is a T3SS structural component (57). *MxiM* deletion mutants do not assemble a functional T3SS, do not secrete T3SS effectors, and do not invade tissue-cultured epithelial cells (57–59). Like the plasmidless strain, the Δ*icsA* and Δ*mxiM* strains did not cause disease; none of the rabbits infected with either of these two mutant strains developed diarrhea, succumbed to infection, or had a reduction in body temperature (Fig. 1A & E). Additionally, none of the mutants induced colonic edema or epithelial cell sloughing, pathologic features that characterized WT infection (Fig. 2 and Fig. 6). Collectively, these data indicate that both IcsA and the T3SS are required for *Shigella* pathogenesis in the infant rabbit model.

**Figure 5.**
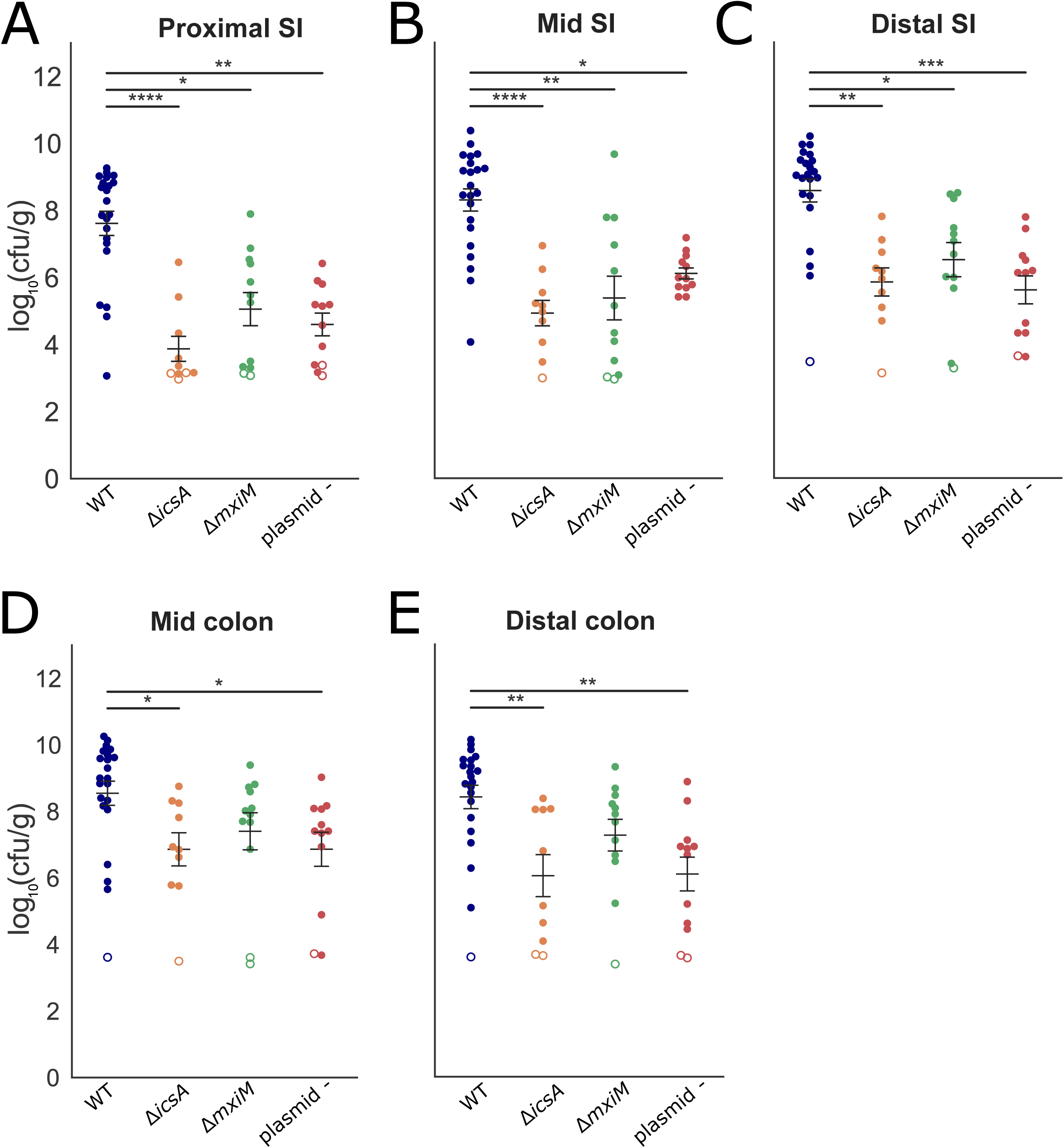
Intestinal colonization of WT and mutant *S. flexneri*. A-E. Bacterial burden of the indicated strains in the indicated intestinal sections 36 hpi. SI = small intestine. Each point represents measurement from one rabbit. Data plotted as log transformed colony forming units (cfu) per gram of tissue. Means and standard error of the mean values are superimposed. Open symbols represent the limit of detection of the assay and are shown for animals where no cfu were recovered. For each section, burdens from all strains were compared to each other; statistical significance was determined using a Kruskal-Wallis test with Dunn’s multiple comparison. P-values: <0.05, *; <0.01, **; <0.001, ***; <0.0001, ****.

**Figure 6.**
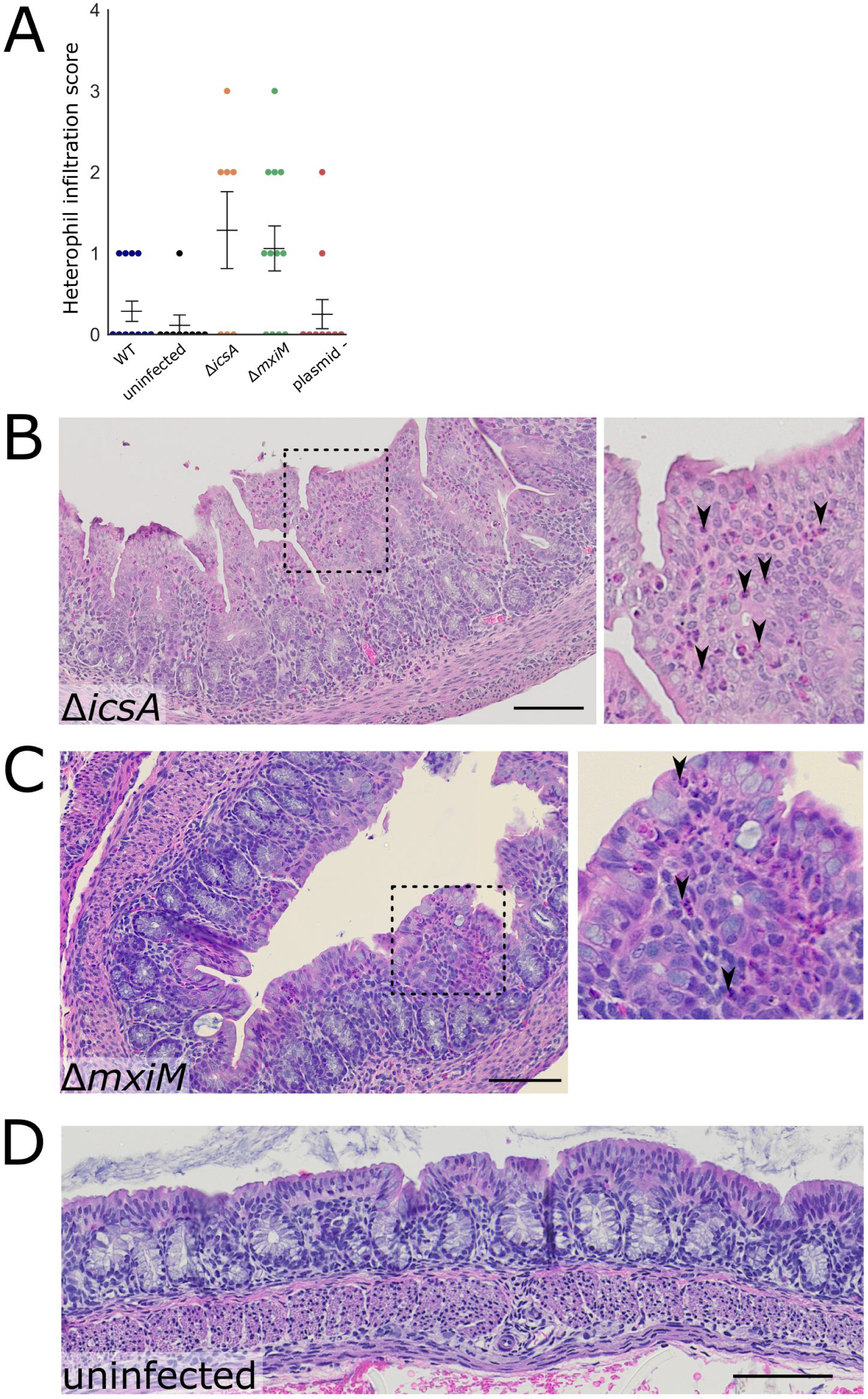
Colonic pathology in rabbits infected with WT or mutant *S. flexneri*. A. Histopathological scores of heterophil infiltration in colonic sections of animals infected with indicated strains of *S. flexneri*. Means and standard error of the mean values are superimposed. Statistical significance was determined using a Kruskal-Wallis test with Dunn’s multiple comparison; comparisons that are non-significant are not labeled. B-D. Representative haematoxylin and eosin-stained colonic sections from rabbits infected with the indicated strains 36 hpi. In B, the inset on the right displays the magnified version of the boxed region of the larger micrograph. Arrowheads point to heterophils (pink cytoplasm, multi-lobular darkly stained nucleus). Scale bar is 100 μm. In C (MxiM mutant), the inset on the right displays a magnified version of the boxed region of the larger micrograph. Arrowheads point to heterophils. Scale bar is 100 μm.

All three of the mutant strains had reduced capacities to colonize the infant rabbit intestine (Fig. 5). Notably, the reduction in the colonization of the *icsA* mutant was at least as great as the other two mutant strains, suggesting that cell-to-cell spreading or the adhesin function of IcsA is critical for intestinal colonization. The colonization defects were most pronounced in the small intestine, where up to 10^4^-fold reductions in recoverable *S. flexneri* cfu were observed (Fig. 5). Reductions in the colon were less marked and did not reach statistical significance for the Δ*mxiM* strain (Fig. 5).

Interestingly, the *icsA* mutant led to an accumulation of heterophils (innate immune cells that are the rabbit equivalent of neutrophils) in the colon that was not observed in animals infected with the WT strain (Fig. 6). Thus, IcsA may contribute to immune evasion by limiting the recruitment of innate immune cells. The *mxiM* mutant also recruited more heterophils to the lamina propria and epithelial cell layer than the WT strain (Fig. 6A & C). Unlike the Δ*icsA* and Δ*mxiM* strains, the plasmidless strain did not recruit heterophils in the colon. Thus, both IcsA and the T3SS appear to antagonize heterophil recruitment, perhaps by facilitating pathogen invasion. However, the absence of heterophil influx in the plasmidless strain challenges this hypothesis and suggests that another plasmid-encoded factor can counteract the actions of IcsA and/or the T3SS in blocking heterophil infiltration.

Since colonic pathology was altered in the mutant strains, we investigated the intestinal localization and IL-8 production induced by the mutants. All three of the mutant strains were found almost exclusively in the lumen of the colon (Fig. 7A, Fig. S4); in contrast to the WT strain (Fig. 3), it was difficult to detect infection foci in the epithelial cell layer in animals infected with mutant strains (Fig. 7). The *icsA* mutant was occasionally observed inside epithelial cells (Fig 7B), but larger foci were not detected. As expected, we observed very few cells expressing IL-8 mRNA in the colons of rabbits infected with any of the three mutant *S. flexneri* strains (Fig. 4D & 7B), supporting the idea that induction of IL-8 expression requires *S. flexneri* invasion of the epithelial cell layer in this model.

**Figure 7.**
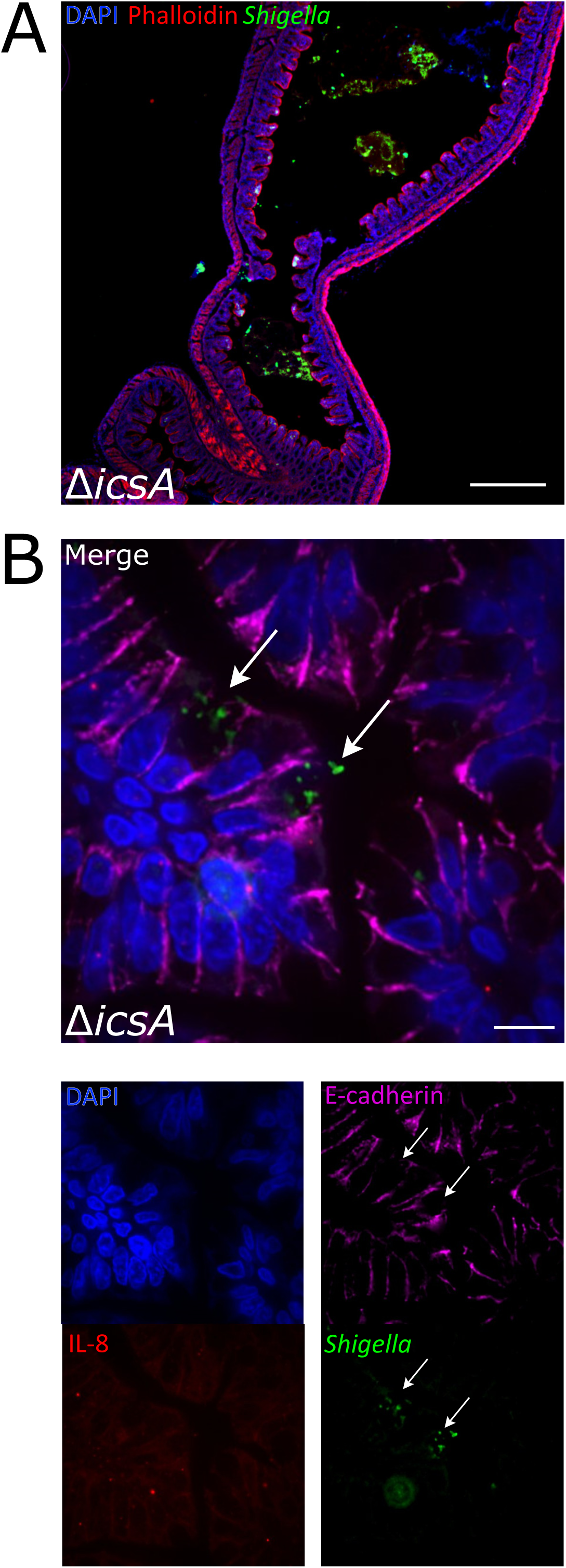
Intestinal localization and IL-8 transcripts in colons from animals infected with an *icsA* mutant. A. Immunofluorescence micrograph of Δ*icsA* in colonic tissue of infected rabbits 36 hpi. Blue, DAPI; green, FITC-conjugated anti-*Shigella* antibody; red, phalloidin-Alexa 568. Scale bar is 500 μm. B. Immunofluorescence micrograph of sections stained with a RNAscope probe to rabbit IL-8 (red), antibodies to *Shigella* (green) and E-cadherin (magenta), and DAPI (blue). Panels on right depicts channels of merged left image. Arrows point to multiple *icsA* bacteria in the cytoplasm of two infected cells. Scale bar is 10 μm.

## Discussion

Small animal models of shigellosis that rely on the oral route of infection have been lacking. Here, we found that oro-gastric inoculation of two to three-day-old infant rabbits with *S. flexneri* led to a diarrheal disease and colonic pathology reminiscent of some aspects of human disease. Fasting animals prior to inoculation reduced the variability in infection outcomes, but not all inoculated animals developed disease. The pathogen robustly colonized the colon, where the organism was found primarily in the lumen; however, prominent infection foci were also observed within the colonic epithelium. Robust *S. flexneri* intestinal colonization, invasion of the colonic epithelium and colonic epithelial sloughing required IcsA and the T3SS, which are both canonical *S. flexneri* virulence factors. Despite the reduced intestinal colonization of the *icsA* and *mxiM* mutants, these strains elicited more pronounced colonic inflammation (characterized by infiltration of heterophils) than the WT strain. IL-8 expression, detected with in situ mRNA labeling, was higher in animals infected with the WT versus the mutant strains, suggesting that epithelial invasion promotes expression of this chemokine. Interestingly, IL-8 expression was greater in uninfected cells near infected epithelial cells than in infected epithelial cells themselves. Collectively, our findings suggest that oral infection of infant rabbits offers a useful experimental model for investigations of the pathogenesis of shigellosis.

Fasted animals developed disease more frequently and had elevated intestinal colonization compared to animals who fed ad libidum prior to inoculation. The presence of inhibitory substances in milk, such as lactoferrin, which degrades components of the *Shigella* T3SS apparatus (39), may limit bacterial establishment in the intestine, but have less potent effects once colonization is established. Mean colonic colonization was higher in animals that developed disease than those that did not (Fig S1A). However, high bacterial burdens are not the only factor predictive of disease; several animals with high pathogen burdens did not exhibit signs of disease (Fig. S1A). Also, initial rabbit body weights did not strongly influence clinical outcomes (Fig. S1C). Several additional factors likely modulate *Shigella* colonization and disease manifestation in infant rabbits. For example, variations in the intestinal microbiota of the infant rabbits may limit or potentiate *S. flexneri* virulence and/or colonization, as described for infections caused by other enteric pathogens, including *Clostridium difficile* (60), *L. monocytogenes* (61), and *V. cholerae* (62). Differences in dam feeding patterns also likely influence colonization and disease outcomes. Further elucidation of factors that modulate outcomes will be valuable to improve this model because they may point to ways to elevate the fraction of animals that develop clinical signs of infection.

A high inoculum dose (10^9^ cfu) was required to achieve reliable disease development following oral inoculation of two to three-day-old infant rabbits. Animals inoculated with lower doses (e.g., 10^7^ cfu) of *S*. *flexneri* developed disease and robust intestinal colonization at lower frequencies. Interestingly, even in oral non-human primate models, the standard inoculum dose (10^10^ cfu) to ensure consistent development of disease (63, 64) is orders of magnitude greater than the dose used in human challenge studies (typically 10^3^-10^4^ cfu) (37, 65, 66). The reasons accounting for these marked differences in infectious doses warrant further exploration. It is unlikely that older rabbits infected via the oral route will be susceptible to colonization and disease, since our findings with other pathogens (35) suggest that infant rabbits become resistant to oral inoculation with enteric pathogens when they are ∼5 days old.

In human infections, *Shigella* causes colonic pathology characterized by an acute inflammatory response with mucosal ulceration and erosions, neutrophil infiltration, congestion, and hemorrhage (8, 9). In the oral infant rabbit model, the WT strain caused edema and sloughing of epithelial cells in the colon, but we did not observe recruitment of heterophils, suggesting that colonic pathology is not primarily attributable to an acute inflammatory response characterized by heterophil infiltration. Instead, the pathology may be driven by invasion and replication of the pathogen within colonic epithelial cells. Oro-gastric inoculation of infant rabbits with EHEC induces heterophil infiltration in the colon (67), indicating that these animals are capable of mounting an acute inflammatory response in this organ.

The marked colonization defect of the Δ*icsA* strain, matching that observed for the Δ*mxiM* (T3SS -) and plasmidless strains, was unexpected. It seems unlikely the Δ*icsA* mutant’s colonization defect is entirely attributable to the mutant’s deficiency in cell-to-cell spreading. Zumsteg et al. found that IcsA can also serve as an adhesin (68). Since distinct regions of IcsA are required for its adhesive versus cell spreading activities (68), it may be possible to genetically dissect which of these IcsA functions plays a dominant role in colonization, using *S. flexneri* strains producing mutant versions of IcsA. Passage of the pathogen through the upper gastrointestinal tract may be required to reveal IcsA’s adhesive activity, because a Δ*icsA* strain had only a modest colonization defect after intra-rectal instillation (10). It was also surprising that the animals infected with the Δ*icsA* strain recruited heterophils to the colon despite little induction of IL-8 expression. These observations suggest that there are additional factors contributing to heterophil recruitment to the rabbit colon. Moreover, since there is minimal heterophil recruitment in animals infected with the WT strain, IcsA-mediated pathogen adherence to colonic epithelial cells (and potentially concomitant increased invasion) may increase delivery of T3SS effectors into host cells, thereby repressing a host-derived heterophil recruitment factor.

Our attempts to utilize TIS to identify novel genetic loci contributing to *S. flexneri* colonization in the infant rabbit intestine were stymied by a narrow infection bottleneck. The tight bottleneck leads to large, random losses of genetic diversity of the input library. The underlying causes of in vivo bottlenecks vary and may include stomach acidity, host innate immune defenses, such as antimicrobial peptides, the number of available niches in the intestine, and competition with the endogenous commensal microbiota (69). Modifications to either the inoculation protocol or library generation could facilitate future in vivo TIS screens. For example, the diversity of the inoculum could be reduced by generating a defined library of transposon mutants with only one or two mutants per gene (e.g. as has been done in *Edwardsiella piscicida* (70)). Regarding the infection protocol, it is possible that the fraction of the inoculum that initially seeds and colonizes the intestine could be elevated by reducing the number of commensal organisms in the intestine that may compete for a niche similar to that occupied by *S. flexneri*. Similar strategies have been utilized to facilitate studies of other enteric pathogens (71, 72).

The intra-rectal infant rabbit model of shigellosis reported by Yum et al. has some beneficial features compared to the oral infection model. Using this route, Yum et al. reported that all animals developed bloody diarrhea and colonic pathology that included substantial recruitment of heterophils (10). As noted above, for unknown reasons, oral inoculation of WT *S. flexneri* did not lead to heterophil recruitment to sites of damage in the colon. An additional difference is that intra-rectal instillation of a Δ*icsA* mutant led to induction of cytokine expression, heterophil recruitment, and only slightly reduced colonization of the strain, whereas following oral inoculation, a Δ*icsA S. flexneri* exhibited a marked colonization defect and did not induce IL-8 mRNA expression. Additional studies are required to elucidate the reasons that account for the differential importance of IcsA in these models. While some features of the intra-rectal model are attractive, Yum et al. used 2 week old rabbits that were carefully hand reared in an animal facility from birth using a complex protocol that may prove difficult for others to adopt (10). In addition to the physiologic route of infection, the oral infant rabbit model requires far less specialized animal husbandry than the intra-rectal model and may therefore prove more accessible.

In summary, oral inoculation of infant rabbits with *Shigella* provides a feasible small animal model to study the pathogenesis of this globally important enteric pathogen. The model should also be useful to test new therapeutics for shigellosis, an issue of increasing importance given the development of *Shigella* strains with increasing resistance to multiple antibiotics (73–76).

## Acknowledgements

This study was supported by the NIGMS grant T32GM007753 (J.D.D.), NIAID grant T32AI-132120 (J.D.D. & A.R.W.), and NIAID grant R01-AI-043247 and the Howard Hughes Medical Institute (M.K.W.).

We gratefully acknowledge Marcia Goldberg for providing *S. flexneri* 2a strains 2457T and BS103 (the virulence plasmidless derivative), and for transducing the streptomycin resistance allele into BS103. We thank Angelina Winbush for help with construction of the Δ*icsA* mutant strain. We thank the Dana-Farber/Harvard Cancer Center in Boston, MA, for the use of the Rodent Histopathology Core, which provided tissue embedding, sectioning, and staining service (NIH 5 P30 CA06516). We thank Rod Bronson at the Rodent Histopathology Core for providing blinded pathology scoring of tissue sections. We thank Brigid Davis and members of the Waldor lab for comments on the manuscript.

## Materials and Methods

### Bacterial strains and growth

Bacterial strains are listed in Table S2. *S*. *flexneri* were routinely grown aerobically in Miller lysogeny broth (LB) or LB agar at 30°C or 37°C. Antibiotics, when used, were included at the following concentrations: Streptomycin (Sm) 200 µg/mL, Kanamycin (Km) 50 µg/mL, Carbenicillin (Carb) 100 µg/mL, Chloramphenicol (Cm) 10 µg/mL. To check for the presence of the virulence plasmid, bacteria were grown on media with Congo red added at 0.1% w/v.

*E. coli* were routinely grown in LB media or agar. Antibiotics were used at the same concentrations as *S. flexneri* except for Cm, which was 30 µg/mL. When required, diaminopimelic acid (DAP) was added at a concentration of 0.3 mM.

### Strain construction

*S. flexneri* 2a strain 2457T and BS103 (a derivative lacking the virulence plasmid) were gifts of Marcia Goldberg. A spontaneous streptomycin resistant strain of *S. flexneri* 2a strain 2457T was generated by plating overnight LB cultures of *S. flexneri* 2a 2457T on 1000 µg/mL Sm LB plates and identifying Sm resistant (Sm^R^) strains that grew as well as the parent strain. The Sm^R^ strain was used as the wild type strain for all subsequent experiments, including animal experiments and construction of mutant strains. Primers (Table S2) were used to amplify the *rpsL* gene in the strain and Sanger sequencing was performed to determine the nature of the mutation resulting in streptomycin resistance. The streptomycin resistance allele was transferred from the Sm^R^ wild type strain into strain BS103 by P1 transduction, yielding a Sm^R^ plasmidless (plasmid -) strain.

Single gene deletion mutants were generated in the WT Sm^R^ strain using the lambda red recombination method, as previously described (77). Resistance cassettes used in the process were amplified from pKD3 (Cm). Mutations generated by lambda red were moved into a clean genetic background by transferring the mutation to the Sm^R^ wild type strain via P1 transduction. Subsequently, antibiotic resistance cassettes were removed via FLP-mediated recombination using pCP20. Retention of the virulence plasmid throughout P1 transduction of the mutation into the parental WT Sm^R^ strain was monitored by plating bacterial mutants on LB + Congo Red to identify red colonies, and by performing multiplex PCR for various genes spread across the virulence plasmid – primers are listed in Table S2.

### Animal Experiments

Rabbit experiments were conducted according to the recommendations of the National Institutes of Health Guide for the Care and Use of Laboratory Animals, the Animal Welfare Act of the United States Department of Agriculture, and the Brigham and Women’s Hospital Committee on Animals, as outlined in Institutional Animal Care and Use Compliance protocol #2016N000334 and Animal Welfare Assurance of Compliance number A4752-01.

Litters of two to three-day-old New Zealand White infant rabbits with lactating adult female (dam) obtained from a commercial breeder (Charles River, Canada or Pine Acres Rabbitry Farm & Research Facility, Norton, MA) were used for animal experiments.

Infant rabbits were administered a subcutaneous injection of Zantac (ranitidine hydrochloride, 50 mg/kg; GlaxoSmithKline) 3 hours prior to inoculation with the wild type (Sm^R^) or isogenic mutants. We attempted to utilize a bicarbonate solution to administer bacteria, but found that *S. flexneri* do not survive when re-suspended in a sodium bicarbonate solution. For initial experiments, a day after arrival, infant rabbits were oro-gastrically inoculated with 1e9 cfu of log phase *S. flexneri* suspended in LB. To prepare the inoculum, an overnight bacterial culture grown at 30°C was diluted 1:100 and grown at 37°C for 3 hours. The bacteria were subsequently pelleted and re-suspended in fresh LB to a final concentration of 2e9 cfu/mL. Rabbits were oro-gastrically inoculated using a PE50 catheter (Becton Dickson) with 0.5 mL of inoculum (1e9 cfu total). In later experiments, infant rabbits were first separated from the dam for 24 hours prior to inoculation, after which they were immediately returned to the dam for the remainder of the experiment.

The infant rabbits were then observed for 36-40 hours post-inoculation and then euthanized via isoflurane inhalation and subsequent intracardiac injection of 6 mEq KCl at the end of the experiment or when they became moribund. Animals were checked for signs of disease every 10-12 hours. Body weight and body temperature measurements were made 1-2 times daily until the end of the experiment. Body temperature was measured with a digital temporal thermometer (Exergen) and assessed on the infant rabbit chest, in between the front legs. Temperatures reported in Figure 1E are the final temperatures prior to euthanasia and change in body weight in Figure 1F is a comparison of the final to initial body weight. Diarrhea was scored as follows: no diarrhea (solid feces, no adherent stool on hindpaw region) or diarrhea (liquid fecal material adhering to hindpaw region). Animal experiments with isogenic mutants were always conducted with litter-mate controls infected with the WT Sm^R^ strain to control litter variation.

At necropsy, the intestine from the duodenum to rectum was dissected, and divided into separate anatomical sections (small intestine, colon) as previously described (54, 78). 1-2 cm pieces of each anatomical section were used for measurements of tissue bacterial burden. Tissue samples were placed in 1x phosphate buffered saline (PBS) with 2 stainless steel beads and homogenized with a bead beater (BioSpec Products Inc.). Serial dilutions were made using 1xPBS and plated on LB+Sm media for enumeration of bacterial cfu. For processing of tissue for microscopy, 1-2 cm pieces of the tissue adjacent to the piece taken for enumeration of bacterial cfu were submerged in either 4% paraformaldehyde (PFA) for frozen sections or 10% neutral-sbuffered formalin (NBF) for paraffin sections.

For gentamicin tissue assays, a 1-2 cm portion of the colon was cut open longitudinally and washed in 1X PBS to remove luminal contents and then incubated in 1mL of 1xDMEM with 100 μg/mL gentamicin for 1 hour at room temperature. The tissue was subsequently washed 3x with 20x volumes of 1x PBS for 30 min with shaking. The tissue was then homogenized and serial dilutions were plated on LB+Sm media for enumeration of bacterial burden.

For Tn-seq experiments, aliquots of the transposon library were thawed and aerobically cultured in LB for 3 hours. The bacteria were pelleted and resuspended in fresh LB to a final concentration of 1e9 cfu per 0.5 mL inoculum. A sample of the input library (1e10 cfu) was plated on a large LB+Sm+Km plate (245 cm^2^; Corning). Bacterial burdens in infected rabbit tissues were determined by plating serial dilutions on LB+Sm+Km plates. The entire colon was homogenized and plated onto a large LB+Sm+Km plate to recover transposon mutants that survived in the colon. Bacteria on large plates were grown for ∼20-22 hours at 30°C, scraped off with ∼10 mL fresh LB, and ∼ 1 mL aliquots were pelleted. The pellets were frozen at −80°C prior to genomic DNA extraction for Tn-seq library construction.

Data from animal experiments were analyzed in Prism (ver. 8; GraphPad). The Mann-Whitney U test or the Kruskal-Wallis test with Dunn’s post-test for multiple comparisons were used to compare the tissue bacterial burdens. A Fisher’s exact test was used to compare the proportion of rabbits that developed diarrhea after infection with various bacterial strains.

### Immunofluorescence Microscopy

Immunofluorescence images were analyzed from 20 wild-type and at least 4 rabbits infected with each of the various mutant bacterial strains, or uninfected rabbits; 2-3 colon sections per rabbit were examined. Tissue samples used for immunofluorescence were fixed in 4% PFA, and subsequently stored in 30% sucrose prior to embedding in a 1:2.5 mixture of OCT (Tissue-Tek) to 30% sucrose and stored at −80°C, as previously described (35). Frozen sections were made at 10 μm using a cryotome (Leica CM1860UV). Sections were first blocked with 5% bovine serum albumin (BSA) in PBS for 1 hour. Sections were stained overnight at 4°C with a primary antibody, diluted in PBS with 0.5% BSA and 0.5% Triton X-100: anti-*Shigella*-FITC (1/1000; #0903, Virostat); anti-E-cadherin 1:100 (610181, BD Biosciences). After washing with 1xPBS - 0.5% Tween20 (PBST), sections were incubated with 647 phalloidin (1/1000; Invitrogen) for 1 hour at room temperature, washed and stained for 5 min with 4′,6-diamidino-2-phenylindole (DAPI) at 2 μg/mL for 5 min, and covered with ProLong Diamond or Glass Antifade (Invitrogen) mounting media. Slides were imaged using a Nikon Ti Eclipse equipped with a spinning disk confocal scanner unit (Yokogawa CSU-Xu1) and EMCCD (Andor iXon3) camera, or with a sCMOS camera (Andor Zyla) for widefield microscopy.

### Histopathology

Tissue samples used for histopathology analysis were fixed in 10% NBF and subsequently stored in 70% ethanol prior to being embedded in paraffin, as previously described (36). Formalin fixed, paraffin embedded (FFPE) sections were made at a thickness of 5 μm. Sections were stained with hematoxylin and eosin (H&E). Slides were assessed for various measures of pathology, e.g. heterophil infiltration, edema, epithelial sloughing, hemorrhage, by a pathologist blinded to the tissue origin. Semi-quantitative scoring for heterophil infiltration were as follows: 0, no heterophils observed; 1, rare heterophils; 2, few heterophils; 3, many heterophils; 4, abundant heterophils. Brightfield micrographs were collected using an Olympus VS120.

### In situ RNA hybridization

Freshly cut FFPE sections (5 μm) were made of the indicated anatomical sections and stored with desiccants at 4°C. Subsequently, sections were processed and analyzed using the RNAscope Multiplex Fluorescent v2 Assay (Advanced Cell Diagnostics USA-ACDbio) combined with immunofluorescence. Briefly, sections were processed following ACDbio recommendations for FFPE sample preparation and pretreatment using 15-minute target retrieval and 25-minute Protease Plus digestion using the RNAscope HybEZ oven for all incubations. An RNAscope C1 probe (OcIL8) to rabbit CXCL8 was developed and used to stain intestinal sections for CXCL8 mRNA expression. C1 probe was detected with Opal 570 dye (Akoya Biosciences) diluted 1:1000 in Multiplex TSA buffer (ACDbio). Sections were also stained with DAPI (2 μg/mL), anti-*Shigella* FITC (1/1000, Virostat), and anti-mouse E-cadherin (1/100; #610181, BDbiosciences). Slides were imaged using a Nikon Ti Eclipse equipped with a spinning disk confocal scanner unit (Yokogawa CSU-Xu1) and EMCCD (Andor iXon3) camera for high magnification images. Slides were imaged using a widefield Zeiss Axioplan 2 microscope through the MetaMorph imaging system for RNAscope signal quantification.

### Quantitative Image Analysis

Images of mid colon tissue sections stained with RNAscope OcIL8, DAPI, and FITC-conjugated anti-*Shigella* antibody were acquired and analyzed using the MetaMorph (7.1.4.0) application. Briefly, tiled 10x images covering the entire length of the tissue section were collected using Multi-Dimensional Acquisition. For analysis of the percentage of IL-8 mRNA expressing cells that were adjacent to infected cells, we analyzed 86 foci of infection at 100x magnification. Exclusive threshold values were set for the DAPI channel or the rhodamine channel independently and applied to all images in the data set. The threshold values for DAPI or rhodamine were used to create a binary mask of each image. The total area under the binary mask was recorded and used to calculate the percent of total tissue (DAPI area under mask) expressing CXCL8 mRNA (rhodamine area under mask) by dividing the values for rhodamine area by the DAPI area for each image. Percentages were graphed using Prism version 8 (GraphPad).

### Transposon library construction and analysis

A transposon library was constructed in *S. flexneri* 2a 2457T Sm^R^ (WT Sm^R^) using pSC189 (79), using previously described protocols (54, 80) with additional modifications. Briefly, *E. coli* strain MFD*pir* (81) was transformed with pSC189. Conjugation was performed between WT Sm^R^ and MFD*pir* pSC189. Overnight LB cultures of WT Sm^R^ (grown at 30°C) and MFD*pir* (grown at 37°C) were mixed and spotted onto 0.45 μm filters on LB+DAP agar plates. The conjugation reaction was allowed to proceed for 2 hours at 30°C. Subsequently, the bacterial mixtures were resuspended in LB and spread across four 245 cm^2^ LB+Sm+Km square plates to generate single separate colonies for a transposon library. The square plates were grown at 30°C for 20 hours. The colonies that formed (∼800,000 total) were washed off with LB (8 mL per plate) and the bacteria from two plates were combined. Two separate 1 mL aliquots of the two combined mixtures were used to start two 1000 mL LB+Km liquid cultures. The cultures were grown aerobically at 30°C with shaking for 3 hours. For each flask, the bacteria were pelleted and resuspended in a small amount of LB before being spread across two 245 cm^2^ LB+Sm+Km square plates and grown at 30°C for 20 hours. The resulting bacteria on the plate were washed off with LB and resuspended. The OD was adjusted to 10 with LB and glycerol so that the final concentration of glycerol was 25%. 1 mL LB + glycerol aliquots were stored at −80°C for later experiments. In addition, 1 mL aliquots were also pelleted, to generate bacterial pellets to serve as sources of genomic DNA for the initial characterization of the transposon library. The pellets were stored at −80°C prior to genomic DNA extraction for Tn-seq library construction.

Tn-seq library construction and data analysis was performed as previously described (54, 55, 82); briefly, genomic DNA was extracted, transposon junctions were amplified, sequencing was performed on an Illumina MiSeq, and data were analyzed using a modified ARTIST pipeline (54, 55). Sequence reads were mapped onto the *S. flexneri* 2a strain 2457T chromosome (Refseq NC_004741.1) and *S. flexneri* 2a strain 301 virulence plasmid (Refseq NC_004851.1). Reads at each TA site were tallied.

## Supplemental Figure Legends

**Figure S1. Factors influencing development of diarrheal disease in infant rabbits after oro-gastric inoculation of *S. flexneri*.**

A-B. Bacterial burden of *S. flexneri* in the indicated intestinal sections 36 hpi; SI = small intestine. Each point represents measurement from one rabbit. Data plotted as log transformed colony forming units (cfu) per gram of tissue. Means and standard error of the mean values are superimposed. Open circles represent the limit of detection of the assay and are shown for animals where no cfu were recovered. (A) ‘disease +’ refers to animals infected with the WT *S. flexneri* strain that developed disease (diarrhea or became moribund early), ‘disease -’ refers to animals infected with the WT *S. flexneri* strain that did not develop disease. (B) ‘unfasted’ refers to animals fed ad libitum prior to inoculation, ‘fasted’ refers to animals separated from dams for 24 hours prior to inoculation.

C. Initial body weights of infant rabbits inoculated with WT *S. flexneri*, grouped based on whether animal developed disease (diarrhea or early mortality). Means and standard error of the mean values are superimposed.

**Figure S2. Examples of severe colonic pathology in infant rabbits inoculated with *S. flexneri* infection.**

A-B. Haematoxylin and eosin-stained colonic sections of severe hemorrhage in lamina propria and colonic lumen from animals infected with the WT strain at 36 hpi. Arrowheads in (A) indicate either an area of hemorrhage in the lamina propria or hemorrhage spreading to the lumen (inset, A). (B) Hemorrhage and epithelial cell sloughing in colonic lumen. Scale bar is 100 μm.

**Figure S3. Range of IL-8 expression in colons of infected infant rabbits.**

(Top left, plot) Percentage of IL-8 expressing cells in each field of view from colonic tissue sections stained with probe to rabbit IL-8 from individual rabbits infected with the WT strain or from uninfected rabbits. Colored dots correspond to micrographs with similar colored borders. Mean values are indicated with bars.

(Micrographs) Immunofluorescence micrographs of colonic sections from uninfected animals or infant rabbits infected with WT *S. flexneri* strain. Sections were stained with a RNAscope probe to rabbit IL-8 (red), and an antibody to *Shigella* (green), and with DAPI (blue).

**Figure S4. Localization of *S. flexneri* mutants in infected infant rabbits.**

A-C. Immunofluorescence micrographs of *S. flexneri* mutants in colonic tissue of infected rabbits 36 hpi. (A) Inset and white arrows show individual *S. flexneri* Δ*icsA* closely associated with the colonic epithelium. Blue, DAPI; green, FITC-conjugated anti-*Shigella* antibody; red, phalloidin-Alexa 568. Scale bar is 500 μm (A-C).

## Supplemental Tables

**Table S1. Transposon library in *Shigella flexneri* 2a strain 2457T**

**Table S2. Strains, plasmids, and oligos**

